# The vagus nerve is critical for regulation of hypothalamic-pituitary-adrenal axis responses to acute stress

**DOI:** 10.1101/2021.06.03.446790

**Authors:** Bailey N. Keller, Angela E. Snyder, Caitlin R. Coker, Elizabeth A. Aguilar, Mary K. O’Brien, Sarah S. Bingaman, Amy C. Arnold, Andras Hajnal, Yuval Silberman

**Author notes:** Corresponding authors, Corresponding author information, Andras Hajnal, Penn State College of Medicine, Neural and Behavioral Sciences, 500 University Dr, H109, Hershey PA 17033, Yuval Silberman, Penn State College of Medicine, Neural and Behavioral Sciences, 500 University Dr, H109, Hershey PA 17033.

## Abstract

The hypothalamic pituitary adrenal (HPA) axis is a critical regulator of physiologic and psychological responses to acute and chronic stressors. HPA axis function is control by numerous feedback inhibitory mechanisms, disruptions of which can lead to various psychiatric conditions, such as depression, posttraumatic stress disorder, and schizophrenia. Vagus nerve stimulation has been shown to be efficacious in the treatment of in these various mental health issues potentially via modulation of HPA axis function, but the mechanisms by which the vagus nerve may regulate HPA function has not been fully elucidated. In the present studies, we sought to test the hypothesis that the vagus nerve is a critical regulator of HPA function. Neuroendocrine function and neurocircuit changes in corticotropin releasing factor (CRF) neurons in the paraventricular nucleus of the hypothalamus (PVN) were examined following acute stress after subdiaphragmatic left vagotomy (VX) in adult male Sprague-Dawley rats. We found that VX mimics HPA activation seen in sham surgery animals exposed to acute restraint stress, particularly increased plasma corticosterone levels, elevated PVN CRF mRNA, and increased action potential firing of putative CRF neurons in PVN brain slices. Furthermore, VX animals exposed to acute restraint stress showed increased elevations of plasma corticosterone and PVN CRF mRNA which may be due to lack of compensatory PVN GABAergic signaling in response to acute stress. Both Sham/Stress and VX/no stress conditions increased action potential firing in putative PVN CRF neurons, but this effect was not seen in the VX/stress condition, suggesting that not all forms of stress compensation are lost following VX. Overall, these findings suggest that the vagus nerve may play a critical role in regulating HPA axis function via modulation of local PVN neurocircuit activity.

## Introduction

The hypothalamic-pituitary-adrenal (HPA) axis is a critical central regulator of behavior, neuroendocrine, and physiologic response to stress in mammals. HPA axis responses to stress are set by activity of corticotropin releasing factor (CRF) neurons in the medial parvocellular region of the paraventricular nucleus of the hypothalamus (PVN). PVN CRF neurons project to the median eminence and upon stimulation release CRF and other peptides into the hypophysial portal vein ending in the anterior pituitary causing the release of adrenocorticotrophic hormone (ACTH). ACTH is released into the systemic circulation and activates cells in the adrenal cortex to synthesize glucocorticoids (cortisol in humans, corticosterone in rodents, CORT) and mineralocorticoids. These hormones perform many functions including providing negative feedback regulation of PVN CRF neuron excitability to return HPA axis function back to homeostatic levels [for reviews of HPA system see (Herman *et al*, 2016, 2020; Smith and Vale, 2006; Spencer and Deak, 2017)]. PVN CRF neuron excitability is controlled by a balance of excitatory glutamatergic and inhibitory GABAergic neurotransmission that arises from a number of upstream brain regions involved in integrating memory and sensory stimuli via both direct and indirect pathways as well as local neurotransmitter and neuropeptide systems. In addition to these pathways, PVN CRF neuron excitability can also be modulated by other neurotransmitter systems such as serotonin, norepinephrine, acetylcholine, and dopamine (Sunstrum and Inoue, 2019). Disruption of these various neurotransmitter and neuropeptide regulators of PVN CRF neuron excitability has been implicated in a myriad of mental health disorders, such as addiction, depression, schizophrenia, and post-traumatic stress disorder (Cáceda *et al*, 2007; Jiang *et al*, 2019).

Another source of HPA axis regulation is the vagus nerve. The vagus nerve, also known as the tenth cranial nerve, is the longest cranial nerve and provides widespread innervation of many organ systems, muscles, and fat depots. The vagus nerve is a mixed nerve comprised of roughly 80% afferent and 20% efferent fibers. As such, the vagus nerve provides a critical interface between the body and the brain, serving as a key point for interoception (Kenny and Bordoni, 2019; Mandalaneni and Rayi, 2020). Previous research suggests an inverse relationship between vagal function and HPA axis activation by stress such that low vagal tone is associated with increased HPA activity in response to stress (Marca *et al*, 2011) and impaired recovery from stress (Weber *et al*, 2010) observed in clinical studies. Compatible with such role, vagus nerve stimulation has been shown to reduce HPA axis function in animal models (Chen *et al*, 2021) and improve clinical outcomes in treatment resistant depression (Sackeim *et al*, 2020) and epilepsy (Mandalaneni and Rayi, 2020). Vagus nerve stimulation has been investigated as experimental treatments for posttraumatic stress disorder and other psychiatric conditions (Cimpianu *et al*, 2017), autoimmune disorders, and chronic inflammatory disorders(Mandalaneni and Rayi, 2020), disorders that often include HPA axis dysregulation. Together, these findings suggest that the vagus nerve may bidirectionally modulate HPA axis function.

The precise mechanisms of how the vagus nerve modulates HPA axis function, however, remains unknown. In particular, how disruption of vagus nerve activity might alter local neurocircuit function in PVN CRF neurons had not been previously characterized. Therefore, the experiments presented here sought to test the hypothesis that damage to the vagus nerve will result in enhanced physiologic stress-responsiveness in adult male rats due to altered neurotransmission and excitatory activity of PVN CRF neurons. To test this hypothesis, we performed subdiaphragmatic vagotomy and examined plasma CORT and CRF mRNA in the PVN and how these change in response to a subsequent acute stress. We further examined how VX and stress conditions modulate putative PVN CRF neuron activity utilizing whole cell patch clamp electrophysiology. We found that VX reduces the ability of local PVN circuits to modulate GABAergic transmission in response to restraint stress. VX was also associated with higher rates of PVN CRF neuron action potential firing. Overall, our findings demonstrate that the vagus nerve plays a role in mediating negative feedback modulation of PVN CRF neurons.

## Methods

### Animals

Male Sprague Dawley rats (260-450 g), obtained from Charles River Laboratories (Horsham, PA, USA), were used in this study. Animals were group housed (3/group) in mesh floored, stainless steel hanging cages in a temperature-controlled vivarium while maintained on a constant 12:12 h light-dark cycle (lights on at 0700). Animals were maintained on pelleted normal rat chow *ad libitum.* Filtered tap water also was available *ad libitum* throughout the experiments. All animal procedures were approved by the Pennsylvania State University College of Medicine Institutional Animal Care and Use Committee and conformed to National Institutes of Health guidelines for the care and use of laboratory animals. Two cohorts were used. The first cohort was utilized for qPCR and ELISA assays while the second cohort was utilized for electrophysiology experiments. Experimenters were blinded to treatment conditions until raw data analysis was completed.

### Subdiaphragmatic vagotomy

All rats received either the vagotomy (VX) or Sham surgery. Rats were given the antibiotic Ceftriaxone (30 mg/kg SC) and Buprenorphine SR (1 mg/kg SC, for pain control) 15 minutes before the surgical incision. Rats were anesthetized with isoflurane (3-5% for induction, 1.5-3% for maintenance). Depth of isoflurane anesthesia was checked by absence of palpebral reflex and by checking for the absence of a toe-pinch. Surgical site was shaved, cleaned with 3 alternating scrubs of Nolvasan/alcohol. During surgery the rat was placed upon a clean, absorbent pad and temperature is maintained at 37 degrees C by the use of circulating warm water blankets. The animal was placed in dorsal recumbency and a midline abdominal skin incision was made. The lobes of the liver were retracted cranially while the stomach was retracted caudally to expose the animal’s diaphragm. Each branch of the main vagus nerve was carefully isolated from the surrounding connective tissue. In VX but not the Sham rats, a small section was excised from the isolated anterior and posterior gastric branches of the vagal nerve. Closure was done in two layers, muscle and skin. The muscle layer was closed using interrupted 3-0 silk suture and the skin was closed using 4-0 nylon (Ethilon monofilament) suture. Postoperatively, Carprofen (30 mg/kg SC) was administered 3 hours after surgery and every day thereafter for 3 days. Rats were then singly housed and allowed to recover for at least 2 weeks before being further utilized.

### Restraint Stress

Rats were immobilized in a plastic restraint bags (AIMS_™_ Body-Contour_™_ Cone Shape Rodent Restraint Bags and left undisturbed for 1 hr. Following restraint stress, rats were moved back in their home cage for 1 hour then lightly anesthetized with isoflurane and euthanized by rapid decapitation. Trunk blood was collected for plasma corticosterone assay and brain tissue dissected and flash frozen in liquid nitrogen and stored at −80C for PCR assays. Non-stress animals were similarly handled but not placed in the restraint bag.

### Plasma corticosterone assay

Plasma corticosterone ELISA assay was performed according to manufacturer recommendations (R&D Systems, KGE009). Blood was collected in 15 ml conical tubes and allowed to clot for 2 hrs at room temperature before being centrifuged for 20 min at 2000x g. Serum was removed, aliquoted, and stored at −20 C until assay was performed.

### qPCR

The paraventricular nucleus of the hypothalamus was micro-dissected, flash frozen in liquid nitrogen, and stored at −80 C until use. Frozen tissue was homogenized in QIAzol Lysis Reagent using a TissueLyser II, with total RNA extracted using RNAeasy Lipid Tissue Mini kits and QIACube automated processing (Qiagen; Germantown, MD). RNA concentration was measured with a NanoDrop spectrophotometer (ND-1000, Thermo Fisher Scientific; Waltham, MA). cDNA was synthesized from total RNA using a high-capacity cDNA reverse transcription kit (ThermoFisher Scientific; Waltham, MA). Quantitative real-time polymerase chain reaction (qPCR) was performed on a QuantStudio 12K Flex system (Applied Biosystems; Foster City, CA) using rat-specific Taqman gene primers (ThermoFisher Scientific; Waltham MA). The primers used were CRF (Rn01462137_m1) and beta-actin (Rn00667869_m1). Each sample was measured in triplicate with cycle threshold (CT) values normalized to beta-actin. Relative gene expression was determined using the 2^-ΔΔCT^ method.

### Electrophysiology

Rats were lightly anesthetized with isoflurane and euthanized via rapid decapitation. Brains were immediately removed and placed in ice-cold sucrose-based artificial cerebrospinal fluid (ACSF). Two hundred and fifty-micrometer-thick coronal brain slices containing the PVN (−1.4 to −2.12 from Bregma) were then prepared in ice-cold sucrose ACSF using a Leica VT1200s vibroslicer with ceramic blades (part#7550/1/C, batch#1710, Campden Instruments Limited). Sucrose based ACSF contained the following (in mM): 183 sucrose, 20 NaCl, 0.5 KCl, 1 MgCl2, 1.4 NaH2PO4, 2.5 NaHCO3, and 1 glucose. Following dissection, slices were transferred to a holding chamber with Modified ACSF [containing (in mM): 100 sucrose, 60 NaCl, 2.5 KCl, 1.4 NaH2PO4, 1.1 CaCl2, 3.2 MgCl2, 2 MgSO4, 22 NaHCO3, 20 glucose, 1 ascorbic acid] at 28°C for 20-30 min and then moved to a separate holding chamber with oxygenated standard ACSF [containing (in mM): 124 NaCl, 4.4 KCl, 2 CaCl2, 1.2 MgSO4, 1 NaH2PO4, 10 glucose, 26 NaHCO3] at 28°C for at least 30 min prior for use in electrophysiology experiments. For electrophysiology recordings, slices were transferred to the submersed recording chamber where they were constantly perfused with standard ACSF with 50 uM APV (to block NMDA-receptor mediated currents) at a rate of 2mL/min at room temperature and allowed to equilibrate for at least 15 min prior to recordings. Electrophysiology recordings were made using SutterPatch 7/8 software utilizing Sutter Integrated Patch Clamp Amplifier and Data Acquisition System (Sutter Instruments). Pipette solution contained (in mM): 135 K+-gluconate, 5 NaCl, 2 MgCl2, 10 HEPES, 0.6 EGTA, 4 Na-ATP, 0.4 Na-GTP, pH 7.2–7.3, 280–290 mOsmol). Whole-cell, voltage-clamp recordings of AMPA receptor-mediated spontaneous excitatory postsynaptic currents (sEPSCs) were made at −55 mV (reversal potential for GABAA currents) while whole-cell voltage clamp recordings of GABA mediated spontaneous inhibitory postsynaptic currents (sIPSCs) in the same cell were made at +10mV (reversal potential for AMPA currents). Cells were allowed to equilibrate to whole-cell configuration for 3–5 min before recordings began and access resistance was monitored continuously. Those experiments in which the access resistance changed by >20% were not included in the data analyses. Cells were then switched to the current-clamp configuration, allowed to equilibrate to their resting membrane potentials for 3-5 minutes, and membrane voltage changes in response to incremental current steps (ranging from −120 pA to +120 pA of 350 ms in duration) were assessed. Three to five cells were used per slice, with each cell being an individual n per experimental group. Putative CRF neurons were targeted by location within the slice and identified post-hoc by electrophysiologic properties (Hoffman et al., 1991; Luther et al., 2002; Luther & Tasker, 2000; Tasker & Dudek, 1991). Recorded neurons that did not correspond to these predefined parameters were excluded from analysis.

### Statistical analyses

Statistical analyses were performed using Microsoft Excel 2016 and GraphPad Prism 8. GraphPad Prism 8 and Microsoft PowerPoint 2016 were used for figure preparation. Specifically, when comparing across treatment groups, a one-way ANOVA was used followed by Tukey’s post-hoc analysis to determine any significance of specific comparisons between groups. All values throughout the study are presented as mean ± SEM.

### Reagents

All reagents used were purchased from Sigma-Aldrich or Fischer Scientific unless otherwise noted in the text.

## Results

### Vagotomy enhances plasma corticosterone response to acute restraint stress

We sought to test the hypothesis that the vagus nerve is required for regulation of HPA stress responses. To test if the vagus nerve plays a critical role in regulating HPA responses to acute stress, sham and VX rats were exposed to a 1 hr restraint stress and plasma was collected 30 min after stress conclusion for assessment of CORT levels via ELISA. Control animals were similarly handled but did not receive restraint. One-way ANOVA indicated a significant difference between groups (F_(3,33)_=3.48, p<0.05, **Fig 1**) with Tukey’s post hoc analysis showing a significant difference between VX/Stress and Sham/non-stress (NS) rats.

**Fig 1.**
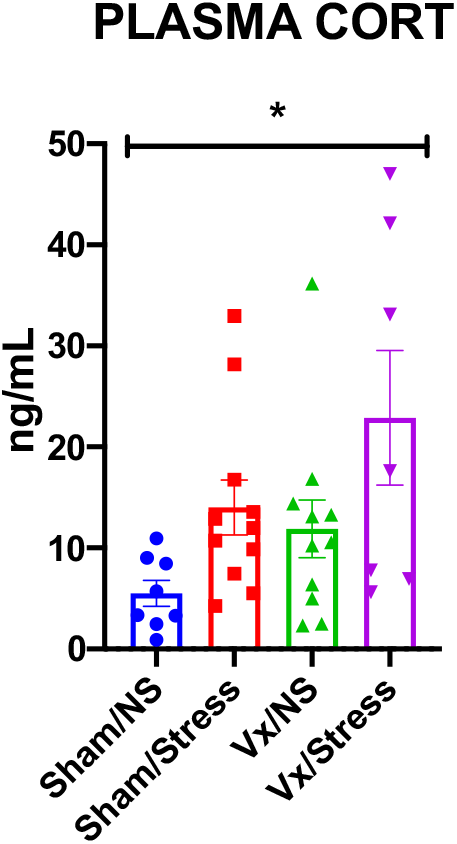
Vagotomy enhances stress-induced corticosterone release. Rats were given either vagotomy (VX) or sham surgery. At least two weeks post-surgical recovery, rats were physically restrained for 1 hr and then plasma CORT collected 1 hr later. One-way ANOVA indicates a significant effect of VX and stress on CORT levels after restraint stress. * indicates p<0.05.

### Vagotomy enhances PVN CRF mRNA response to acute restraint stress

To test if the enhanced stress evoked plasma CORT elevation by VX was due to changes in HPA function, we assessed CRF mRNA in PVN tissue in the animals above. One-way ANOVA indicated a significant difference between groups (F_(3,25)_=5.330, p<0.01, **Fig 2**) with Tukey’s post hoc analysis showing a significant difference between VX/Stress and Sham/NS rats.

**Fig 2.**
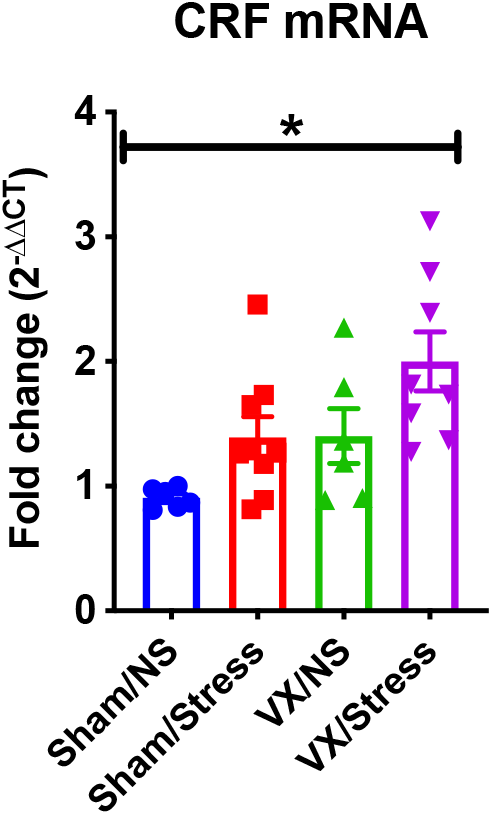
Vagotomy enhances stress-induced CRF mRNA in the PVN. From the same rats were given either vagotomy (VX) or sham surgery as Fig 1. At least two weeks post-surgical recovery, rats were physically restrained for 1 hr and PVN tissue collected 1 hr later. One-way ANOVA indicates a significant effect of VX and stress on PVN CRF mRNA expression after restraint stress. * indicates p<0.05.

### Vagotomy and stress modulate excitatory and inhibitory neurotransmission and action potential firing in putative PVN CRF neurons

We next sought to determine if VX and stress altered excitatory and inhibitory neurotransmission in PVN CRF neurons. Putative PVN CRF neurons were recorded via whole-cell patch clamp and spontaneous excitatory postsynaptic currents (sEPSC) and spontaneous inhibitory postsynaptic currents (sIPSC) were recorded in the same cell by altering the holding potential. There were no significant differences in sEPSC frequency between groups (F_(3,42)_=0.181, p>0.05) but there was a significant difference in sEPSC amplitude (F_(3,42)_=5.582, p<0.01). Post-hoc analysis showed a significant difference between VX/NS group compared to the Sham/Stress and VX/Stress groups (p<0.05). There was a significant difference in sIPSC frequency (F_(3,42)_=2.844, p<0.05) but no significant difference in sIPSC amplitude (F_(3,42)_=0.636, p>0.05) between groups. We then assessed the ratio of excitatory and inhibitory current frequencies (E/I ratio) changes across groups and found a significant difference across groups (F_(3,42)_=5.379, p<0.01). Post hoc analysis showed a significant difference between VX/Stress compared to all other groups (p<0.05).

**Fig 3.**
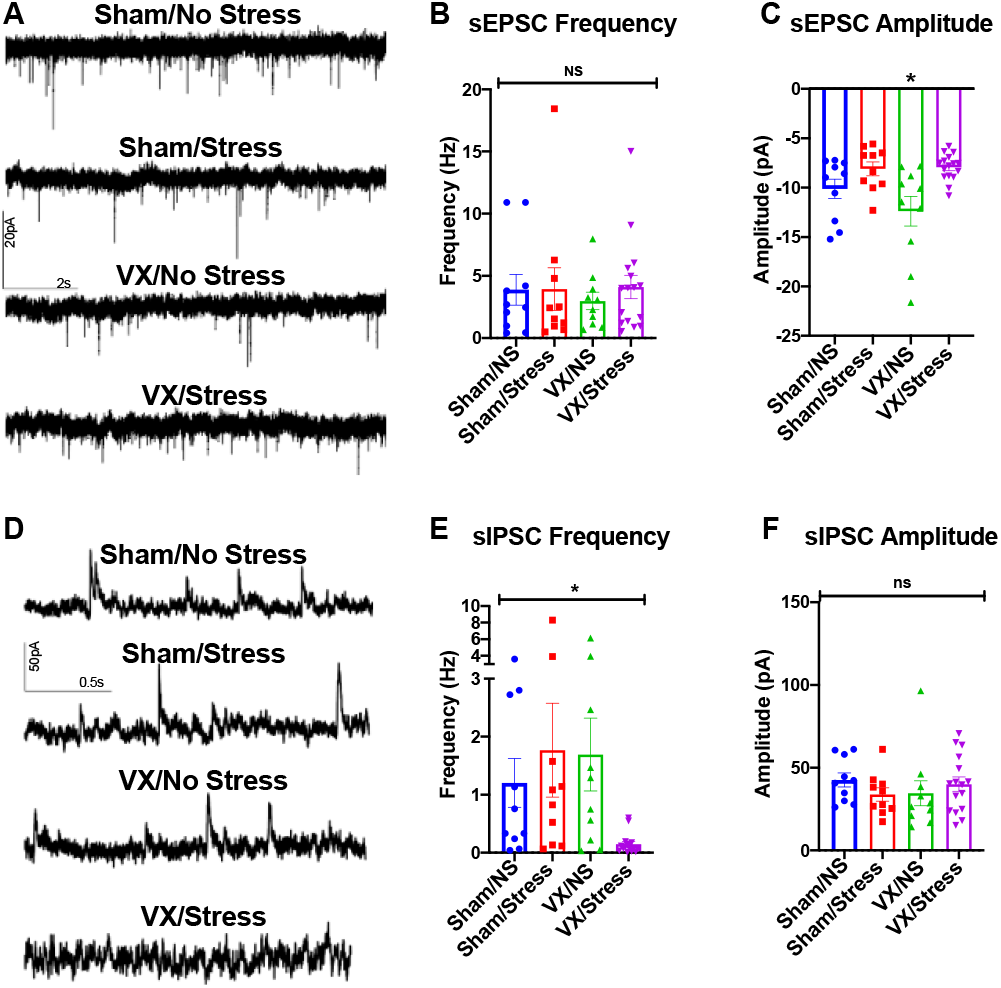
Vagotomy reduces inhibitory neurotransmission in the PVN after stress. Rats were given either vagotomy (VX) or sham surgery. At least two weeks post-surgical recovery, rats were physically restrained for 1 hr and PVN tissue collected 1 hr later. A-C) Spontaneous excitatory postsynaptic currents (sEPSCs) and (D-E) spontaneous inhibitory postsynaptic currents (sIPSCs) were recorded from putative CRF neurons in the PVN. One-way ANOVA indicates no significant effect of VX and stress on sEPSC frequency, but a significant increase sEPSC amplitude only in the vX/NS group. One-way ANOVA indicates a significant effect of VX and stress on sIPSC frequency, with no significant effects of sIPSC amplitude. * indicates p<0.05.

**Fig 4.**
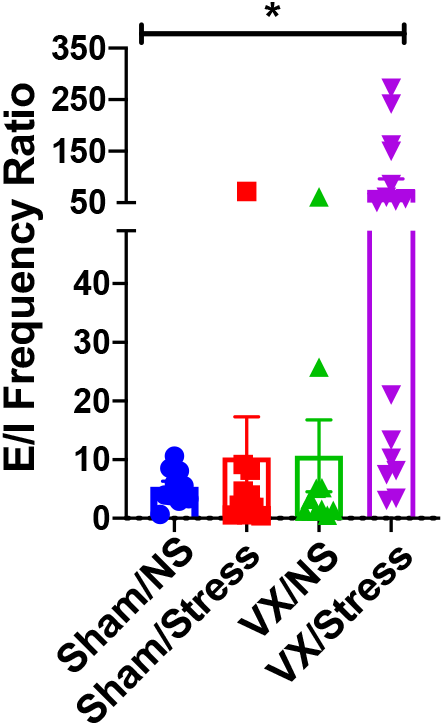
Vagotomy alters excitatory/inhibitory ratio in the PVN after stress. From data shown in Fig 3, excitatory/inhibitory (E/I) ratio was determined by dividing sEPSC frequency by sIPSC frequency. One-way ANOVA indicates a significant effect of VX and stress on E/I ratio. * indicates p<0.05.

In the same neurons as above, we then examined action potential firing rates evoked by different current steps. Two-way ANOVA indicated a significant main effect of groups on action potential firing rates (F_(3,317)_=8.03, p<0.001).

**Fig 5.**
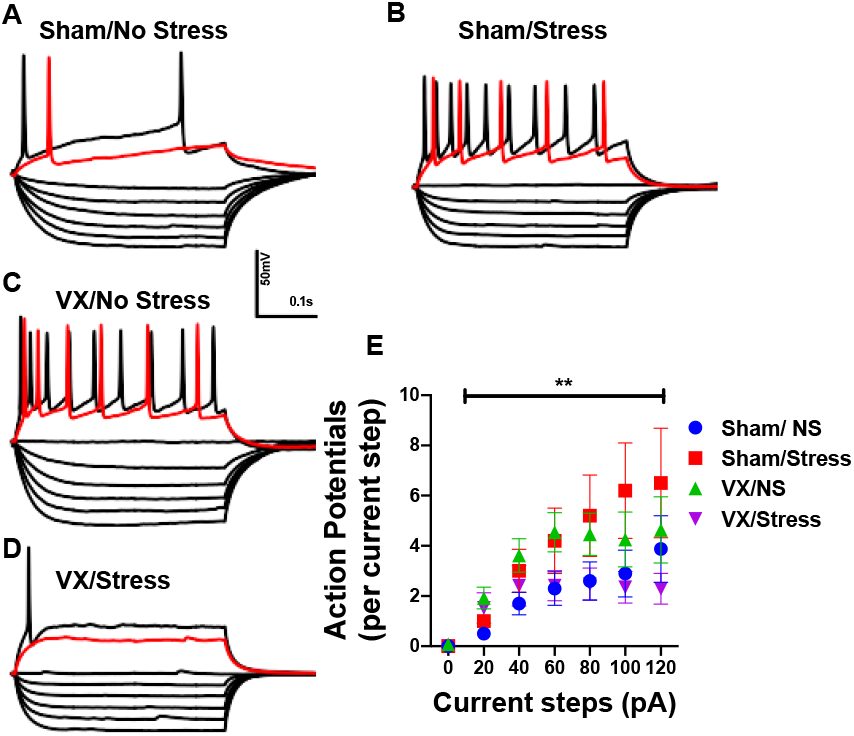
Vagotomy increases putative CRF neuron action potential firing similarly to acute stress. Following recordings of sEPSC and sIPSCs, action potential firing was recorded in the neurons above. A-D) Example recordings of membrane potentials in response to hyperpolarizing and depolarizing currents in the various VX and stress conditions. The positive 60pA step for each neuron in shown in red. E) Summary graph of average number of action potentials per depolarizing step Two-way ANOVA indicates a significant main group effect on action potential firing. ** indicates p<0.01.

## Discussion

Vagus nerve activity is inversely correlated with HPA activity although the mechanisms behind this phenomenon are not clear. Here, we sought to elucidate if the vagus nerve plays a role in resting and stress induced activity of the HPA axis and putative CRF neurons in the PVN by surgically ablating the subdiaphragmatic vagus nerve. Our findings indicate that subdiaphragmatic VX increases basal and stress-induced plasma CORT and PVN CRF mRNA. At the cellular and circuit level, VX induced an increase in putative CRF neuron action potential firing, likely accounting in part for elevated PVN CRF mRNA and plasma CORT. Together, these findings suggest that VX induced a condition similar to a chronic, non-habituating stress exposure. Further, VX increased presynaptic GABAergic transmission, which we propose to be involved in an attempted compensatory mechanism to inhibit over-activated PVN CRF neuron function. VX animals however are unable to mount a compensatory GABAergic response to stress. Overall, these results suggest that the vagus nerve may be a critical component for regulation of PVN CRF neuron activity and HPA axis function following acute stress.

Our results show that stress increases plasma CORT levels in Sham rats, a common occurrence after restraint stress. VX/NS rats had higher basal CORT levels than Sham/NS rats to a level similar to that seen in Sham/stress rats, suggesting VX may induce a state of chronic HPA axis activation. From this elevated basal state, VX/stress rats had higher CORT levels than all other groups, suggesting the chronic stress-like state of VX does not habituate like a repeated homotypic stressor. These findings suggest that VX may induce a stress-like state similar to chronic variable stress or similar heterotypic stress exposures. One limitation of the current study is that CORT was only examined at one time point following a 1 hr restraint. Future studies will need to address this limitation utilizing within subject design with repeated CORT sampling over a longer time frame. Similar to the effects seen with plasma CORT, our data show significant increases in CRF mRNA expression in PVN tissue samples in the VX/Stress group compared to Sham/NS rats. Previous research has shown that most stressors increase PVN CRF mRNA [for review see (Herman and Tasker, 2016)]. Such increased CRF mRNA expression has been shown to result in increased CRF peptide production in the PVN and increased CRF turnover in the median eminence (Chappell *et al*, 1986).

The current findings that VX increases plasma CORT and PVN CRF mRNA are in contrast to previous research showing that subdiaphragmatic VX does not alter stress stimulated CORT in rats (Klarer *et al*, 2014). It should also be noted that the previous study performed both a left subdiaphragmatic VX and a left cervical vagus rhizotomy. This same study also showed that VX reduced anxiety-like behavior in the elevated plus maze and open field and reduced latency to eat in a second trial of a novelty suppressed feeding test. However, these animals also showed reduced ability to extinguish freezing behaviors in an auditory fear conditioning task suggesting a complex role for vagal function in stress-related behaviors both in basal and stimulated conditions. Other studies further suggest that VX induces anhedonia-like behavior in multiple behavioral tasks (Klarer *et al*, 2019). However, previous studies utilizing the same limited VX surgery model used here showed increased alcohol intake (Orellana *et al*, 2021). Overall, future studies will be needed to address these disparities to determine the independent roles of both afferent and efferent vagal fibers in stress-related behaviors and PVN responsively.

The effects of stress on PVN CRF neurocircuit activity and intrinsic excitability are complex and show distinct modulation of glutamatergic and GABAergic input to PVN CRF neurons as well as intrinsic excitability following acute stressors of various types or repeated homotypic vs heterotypic stressors (Herman *et al*, 2002, 2004; Levy and Tasker, 2012). Glutamatergic stimulation increases PVN CRF neuron activity (Feldman and Weidenfeld, 1997). Previous work shows that acute stress rapidly decreases AMPA mediated glutamatergic transmission in the PVN in a glucocorticoid dependent mechanism that may require endocannabinoid signaling (Di *et al*, 2003; Levy and Tasker, 2012; Malcher-Lopes *et al*, 2006; Nahar *et al*, 2015). Chronic stress appears to increase functional AMPA mediated glutamatergic signaling in PVN CRF neurons (Flak *et al*, 2009; Franco *et al*, 2016). Therefore, it is surprising that we saw no change in glutamatergic signaling across the acute stress or VX conditions, except for a small but significant change in sEPSC amplitude in the VX/NS rats. Future work will be needed to examine the discrepancies between the current work and previous studies, but one possibility is the anesthesia from the surgical procedure may have altered glutamate plasticity in this region. Other work has also previously indicated NMDA receptor modulation following acute and chronic stress (Bartanusz *et al*, 1995; Kuzmiski *et al*, 2010; Ziegler *et al*, 2005); however, NMDA receptors were blocked during these recordings so such findings cannot be confirmed in the current studies.

GABAergic receptor activation is thought to reduce PVN CRF neuron activity by both tonic and phasic inhibition (Boudaba *et al*, 1996; Cole and Sawchenko, 2002; Ziegler and Herman, 2002). Although tonic inhibition was not examined in this study, previous research indicates that stress upregulates GABAergic mRNA expression in multiple presynaptic neurons that project to the PVN as part of a compensatory mechanism to return PVN CRF neuron activity to homeostatic levels. For review of GABAergic effects on PVN CRF neuron activity please see (Cullinan *et al*, 2008). In this study, VX/stress animals showed heightened plasma CORT, heightened PVN CRF mRNA expression, and a reduction in GABAergic signaling at the time point tested, suggesting that these animals may not be able to appropriately respond to stressor due to a lack of dynamic GABAergic compensation.

It should be noted, however, that stress can modulate PVN GABAergic signaling in other ways not examined here. In particular, both acute and chronic stress have been shown to alter the chloride gradient in PVN CRF neurons such that GABAergic transmission can lead to GABA-mediated enhancement of neuronal activity as opposed to the typical inhibitory response to GABA (Gao *et al*, 2016; Hewitt *et al*, 2009). Additionally, both acute and chronic stress have also been shown to reduce GABAergic inputs to PVN parvocellular, putative CRF, neurons (Joëls *et al*, 2004; Verkuyl *et al*, 2004); which is likely due to stress induced elevations in glucocorticoids (Herman *et al*, 1995; Miklós and Kovács, 2002; Verkuyl *et al*, 2004). The reason for this apparent discrepancy between studies is unclear, but it might have to do with the timing of recordings. In these studies, rats were restrained for 1 hr and allowed to rest for 1 hr prior to brain slice preparation. It is important to note that all animals had undergone either a VX or Sham surgery, so it is possible that the initial surgical procedures and/or anesthesia may have influenced stress adaptation of PVN GABAergic signals. Overall, the increase in GABAergic signaling following stress may be due to a form of compensation allowing PVN CRF neurons to return to homeostatic activity in the face of elevated plasma CORT. Under these conditions, the lack of enhanced GABAergic signaling in VX/Stress rats in response to further elevations of plasma CORT suggests that the vagus may play a critical role in this potential mechanism of acute stress compensation. This lack of GABAergic signaling change in these rats resulted in a large alteration in E/I ratio not seen in other animals, further supporting the hypothesis that VX results in lack of stress adaptation in these studies. Indeed, the high E/I ratio in the VX/Stress rats is similar to the elevated mEPSC frequency and reduced mIPSC frequency seen in PVN CRF neurons from rats exposed to chronic variable stress, another model of non-adaptive stress (Franco *et al*, 2016).

Our data indicate that acute restraint stress increases action potential firing of putative PVN CRF neurons in brain slices from sham rats. Previous research indicates that acute stress increases PVN CRF neuron activity in vivo although other work has shown minimal changes to PVN CRF neuron action potential firing in brain slices(Kim *et al*, 2019b, 2019a). Supporting our hypothesis that VX-induced a non-habituating stress-like condition, action potential firing in VX rats was also increased compared to Sham/NS rats. Surprisingly, although VX/Stress rats had higher plasma CORT and higher CRF mRNA, action potential firing rates in VX/Stress rats were essentially the same as Sham/NS rats. Together, these findings suggest that VX rats can undergo an acute stress response and allow for some form of compensation in an attempt to return back to homeostatic levels that does not require GABAergic transmission. This potentially represents a novel form of stress compensation dependent on vagus nerve activation, suggesting that the vagus nerve can modulate PVN CRF neuron function via GABAergic signaling and other, currently unknown, mechanisms.

There are a number of potential mechanisms that may account for VX regulation of HPA axis function (Herman *et al*, 2008, 2020; Herman and Tasker, 2016). The first central relay of the vagus nerve is the nucleus of the tractus solitarius (NTS) (Bonaz *et al*, 2018; Kenny and Bordoni, 2019), a medullary region known to send a variety of projections to the PVN (Herman, 2018). These NTS projections contain numerous neurotransmitters and neuropeptides such as glutamate, norepinephrine, and glucagon-like peptide 1 (GLP1). Among these, norepinephrine (NE) has been most widely studied, showing that NE typically increases HPA stress responses, although this may depend on context, adrenergic receptor subtype activation, and type of stress. Chronic stress has also been shown to increase dopamine beta hydroxylase-positive connections with PVN CRF neurons, suggesting NE may play an important role in HPA axis regulation. However, the NTS is not the sole source of PVN NE as the locus coeruleus also sends NE projections to the PVN that may provide a distinct source of stress-stimulated PVN NE signaling (Itoi and Sugimoto, 2010). Additionally, while NE in the PVN is generally thought to be a pro-stress signal via stimulation of CRF neurons, NE can also activate other PVN neurons, such as oxytocin neurons (which increase GABAergic signaling in the PVN), or influence other PVN inputs to reduce stress-responding (Maniscalco and Rinaman, 2018).

In summary, here we show that the vagus nerve is critical in multiple aspects of HPA axis function by modulating both intrinsic PVN CRF neuron excitability and local GABAegric signaling. Although more research is needed to determine the precise mechanisms, these data may help explain, at least in part, how vagus nerve stimulation may improve various psychiatric pathologies that include HPA axis dysfunction.

## Acknowledgements

We thank Nelli Horvath for her assistance with the surgeries and post-operative care. Funding by NIH grants AA026865, AA022937, AA027697, AA027943, TR002014, and TR002016

## References

Bartanusz V, Aubry JM, Pagliusi S, Jezova D, Baffi J, Kiss JZ (1995). Stress-induced changes in messenger RNA levels of N-methyl-d-aspartate and AMPA receptor subunits in selected regions of the rat hippocampus and hypothalamus. Neuroscience 66:247–252.

Bonaz B, Bazin T, Pellissier S (2018). The Vagus Nerve at the Interface of the Microbiota-Gut-Brain Axis. Front Neurosci 12: 49.

Boudaba C, Szabó K, Tasker JG (1996). Physiological mapping of local inhibitory inputs to the hypothalamic paraventricular nucleus. J Neurosci 16:7151–7160.

Cáceda R, Kinkead B, Nemeroff CB (2007). Involvement of Neuropeptide Systems in Schizophrenia: Human Studies. Int Rev Neurobiol 78:327–376.

Chappell PB, Smith MA, Kilts CD, Bissette G, Ritchie J, Anderson C, et al (1986). Alterations in corticoptropin-releasing factor-like immunoreactivity in discrete rat brain regions after acute and chronic stress. J Neurosci 6:2908–2914.

Chen X, Liang H, Hu K, Sun Q, Sun B, Bian L, et al (2021). Vagus nerve stimulation suppresses corticotropin-releasing factor-induced adrenocorticotropic hormone release in rats. Neuroreport 32:792–796.

Cimpianu CL, Strube W, Falkai P, Palm U, Hasan A (2017). Vagus nerve stimulation in psychiatry: a systematic review of the available evidence. J Neural Transm 124:145–158.

Cole RL, Sawchenko PE (2002). Neurotransmitter regulation of cellular activation and neuropeptide gene expression in the paraventricular nucleus of the hypothalamus. J Neurosci 22:959–969.

Cullinan WE, Ziegler DR, Herman JP (2008). Functional role of local GABAergic influences on the HPA axis. Brain Struct Funct 213:63–72.

Di S, Malcher-Lopes R, Halmos KC, Tasker JG (2003). Nongenomic glucocorticoid inhibition via endocannabinoid release in the hypothalamus: A fast feedback mechanism. J Neurosci 23:4850–4857.

Feldman S, Weidenfeld J (1997). Hypothalamic mechanisms mediating glutamate effects on the hypothalamo-pituitary-adrenocortical axis. J Neural Transm 104:633–642.

Flak JN, Ostrander MM, Tasker JG, Herman JP (2009). Chronic Stress-Induced Neurotransmitter Plasticity in the PVN. J Comp Neurol 517:156–165.

Franco AJ, Chen C, Scullen T, Zsombok A, Salahudeen AA, Di S, et al (2016). Sensitization of the Hypothalamic-Pituitary-Adrenal Axis in a Male Rat Chronic Stress Model. Endocrinology 157:2346–2355.

Gao Y, Zhou JJ, Zhu Y, Kosten T, Li DP (2016). Chronic Unpredictable Mild Stress Induces Loss of GABA Inhibition in Corticotrophin-Releasing Hormone-Expressing Neurons through NKCC1 Upregulation. Neuroendocrinology 104:194–208.

Herman JP (2018). Regulation of Hypothalamo-Pituitary-Adrenocortical Responses to Stressors by the Nucleus of the Solitary Tract/Dorsal Vagal Complex. Cell Mol Neurobiol 38:25–35.

Herman JP, Adams D, Prewitt C (1995). Regulatory Changes in Neuroendocrine Stress-Integrative Circuitry Produced by a Variable Stress Paradigm. Neuroendocrinology 61:180–190.

Herman JP, Flak J, Jankord R (2008). Chronic stress plasticity in the hypothalamic paraventricular nucleus. Prog Brain Res 170:353–364.

Herman JP, McKlveen JM, Ghosal S, Kopp B, Wulsin A, Makinson R, et al (2016). Regulation of the hypothalamic-pituitary-adrenocortical stress response. Compr Physiol 6:603–621.

Herman JP, Mueller NK, Figueiredo H (2004). Role of GABA and glutamate circuitry in hypothalamo-pituitary-adrenocortical stress integration. Ann N YAcadSci 1018:35–45.

Herman JP, Nawreen N, Smail MA, Cotella EM (2020). Brain mechanisms of HPA axis regulation: neurocircuitry and feedback in context Richard Kvetnansky lecture. Stress 23:617–632.

Herman JP, Tasker JG (2016). Paraventricular hypothalamic mechanisms of chronic stress adaptation. Front Endocrinol (Lausanne) 7: 137.

Herman JP, Tasker JG, Ziegler DR, Cullinan WE (2002). Local circuit regulation of paraventricular nucleus stress integration: Glutamate-GABA connections. Pharmacol Biochem Behav 71:457–468.

Hewitt SA, Wamsteeker JI, Kurz EU, Bains JS (2009). Altered chloride homeostasis removes synaptic inhibitory constraint of the stress axis. Nat Neurosci 12:438–443.

Hoffman NW, Tasker JG, Dudek FE (1991). Immunohistochemical differentiation of electrophysiologically defined neuronal populations in the region of the rat hypothalamic paraventricular nucleus. J Comp Neurol 307:405–416.

Itoi K, Sugimoto N (2010). The brainstem noradrenergic systems in stress, anxiety and depression. J Neuroendocrinol 22:355–361.

Jiang Z, Rajamanickam S, Justice NJ (2019). CRF signaling between neurons in the paraventricular nucleus of the hypothalamus (PVN) coordinates stress responses. Neurobiol Stress 11:.

Joёls M, Karst H, Alfarez D, Heine VM, Qin Y, Riel E Van, et al (2004). Effects of chronic stress on structure and cell function in rat hippocampus and hypothalamus. Stress 7:221–231.

Kenny BJ, Bordoni B (StatPearls Publishing: 2019). Neuroanatomy, Cranial Nerve 10 (Vagus Nerve). StatPearls at <http://www.ncbi.nlm.nih.gov/pubmed/30725856>.

Kim J, Lee S, Fang YY, Shin A, Park S, Hashikawa K, et al (2019a). Rapid, biphasic CRF neuronal responses encode positive and negative valence. Nat Neurosci 22:576–585.

Kim JS, Han SY, Iremonger KJ (2019b). Stress experience and hormone feedback tune distinct components of hypothalamic CRH neuron activity. Nat Commun 10:.

Klarer M, Arnold M, Günther L, Winter C, Langhans W, Meyer U (2014). Gut vagal afferents differentially modulate innate anxiety and learned fear. J Neurosci 34:7067–7076.

Klarer M, Weber-Stadlbauer U, Arnold M, Langhans W, Meyer U (2019). Abdominal vagal deafferentation alters affective behaviors in rats. J Affect Disord 252:404–412.

Kuzmiski JB, Marty V, Baimoukhametova D V., Bains JS (2010). Stress-induced priming of glutamate synapses unmasks associative short-term plasticity. Nat Neurosci 13:1257–1264.

Levy BH, Tasker JG (2012). Synaptic regulation of the hypothalamic-pituitary-adrenal axis and its modulation by glucocorticoids and stress. Front Cell Neurosci 6: 24.

Luther JA, Daftary SS, Boudaba C, Gould GC, Halmos KC, Tasker JG (2002). Neurosecretory and non-neurosecretory parvocellular neurones of the hypothalamic paraventricular nucleus express distinct electrophysiological properties. J Neuroendocrinol 14:929–932.

Luther JA, Tasker JG (2000). Voltage-gated currents distinguish parvocellular from magnocellular neurones in the rat hypothalamic paraventricular nucleus. J Physiol 523:193–209.

Malcher-Lopes R, Di S, Marcheselli VS, Weng FJ, Stuart CT, Bazan NG, et al (2006). Opposing crosstalk between leptin and glucocorticoids rapidly modulates synaptic excitation via endocannabinoid release. J Neurosci 26:6643–6650.

Mandalaneni K, Rayi A (StatPearls Publishing: 2020). Vagus Nerve Stimulator. StatPearls at <http://www.ncbi.nlm.nih.gov/pubmed/32965846>.

Maniscalco JW, Rinaman L (2018). Vagal interoceptive modulation of motivated behavior. Physiology 33:151–167.

Marca R La, Waldvogel P, Thörn H, Tripod M, Wirtz PH, Pruessner JC, et al (2011). Association between Cold Face Test-induced vagal inhibition and cortisol response to acute stress. Psychophysiology 48:420–429.

Miklós IH, Kovács KJ (2002). GABAergic innervation of corticotropin-releasing hormone (CRH)-secreting parvocellular neurons and its plasticity as demonstrated by quantitative immunoelectron microscopy. Neuroscience 113:581–592.

Nahar J, Haam J, Chen C, Jiang Z, Glatzer NR, Muglia LJ, et al (2015). Rapid nongenomic glucocorticoid actions in male mouse hypothalamic neuroendocrine cells are dependent on the nuclear glucocorticoid receptor. Endocrinology 156:2831–2842.

Orellana ER, Nyland JE, Horvath N, Hajnal A (2021). Vagotomy increases alcohol intake in female rats in diet dependent manner: Implications for increased alcohol use disorder after roux-en-y gastric bypass surgery. Physiol Behav 235:.

Sackeim HA, Dibué M, Bunker MT, Rush AJ (2020). The Long and Winding Road of Vagus Nerve Stimulation: Challenges in Developing an Intervention for Difficult-to-Treat Mood Disorders. doi:10.2147/NDT.S286977.

Smith SM, Vale WW (2006). The role of the hypothalamic-pituitary-adrenal axis in neuroendocrine responses to stress. Dialogues Clin Neurosci 8:383–395.

Spencer RL, Deak T (2017). A users guide to HPA axis research. Physiol Behav 178:43–65.

Sunstrum JK, Inoue W (2019). Heterosynaptic modulation in the paraventricular nucleus of the hypothalamus. Neuropharmacology 154:87–95.

Tasker JG, Dudek FE (1991). Electrophysiological properties of neurones in the region of the paraventricular nucleus in slices of rat hypothalamus. J Physiol 434:271–293.

Verkuyl JM, Hemby SE, Jöels M (2004). Chronic stress attenuates GABAergic inhibition and alters gene expression of parvocellular neurons in rat hypothalamus. Eur J Neurosci 20:1665–1673.

Weber CS, Thayer JF, Rudat M, Wirtz PH, Zimmermann-Viehoff F, Thomas A, et al (2010). Low vagal tone is associated with impaired post stress recovery of cardiovascular, endocrine, and immune markers. Eur J Appl Physiol 109:201–211.

Ziegler DR, Cullinan WE, Herman JP (2005). Organization and regulation of paraventricular nucleus glutamate signaling systems: N-methyl-D-aspartate receptors. J Comp Neurol 484:43–56.

Ziegler DR, Herman JP (2002). Neurocircuitry of stress integration: Anatomical pathways regulating the hypothalamo-pituitary-adrenocortical axis of the rat. Integr Comp Biol 42:541–551.

